# Polygonal motion and adaptable phototaxis via flagellar beat switching in *Euglena gracilis*

**DOI:** 10.1101/292896

**Authors:** Alan C. H. Tsang, Amy T. Lam, Ingmar H. Riedel-Kruse

## Abstract

Biological microswimmers exhibit versatile strategies for sensing and navigating their environment ^1–7^, e.g., run-and-tumble ^2^ and curvature modulation ^3^. Here we report a striking behavior of *Euglena gracilis*, where Euglena cells swim in polygonal trajectories due to exposure to increasing light intensities. While smoothly curved trajectories are common for microswimmers ^3, 8^, such quantized ones have not been reported previously. This polygonal behavior emerges from periodic switching between the flagellar beating patterns of helical swimming ^6, 9^ and spinning ^10^ behaviors. We develop and experimentally validate a biophysical model that describes the phase relationship between the eyespot, cell orientation, light detection, and cellular reorientation, that accounts for all three behavioral states. Coordinated switching between these behaviors allows ballistic, superdiffusive, diffusive, or subdiffusive motion ^11,12^ (i.e., the tuning of the diffusion constant over 3 orders of magnitude) and enables navigation in structured light fields, e.g., edge avoidance and gradient descent. This feedback-control links multiple system scales (flagellar beats, cellular behaviors, phototaxis strategies) with implications for other natural and synthetic microswimmers ^13^.

---

Biological microswimmers exhibit a variety of intricate behaviors and strategies in order to achieve navigational tasks in response to environmental stimuli such as chemicals ^1–3^, light ^4–6^, electric fields ^14,15^, and fluid flows ^7^. Experimental and theoretical studies in many microorganisms (e.g., *Escherichia coli* ^2^, *Chlamydomonas reinhardtii* ^16–20^, *Euglena gracilis* ^6, 9, 21, 22^, *Volvox globator* ^4^, *Paramecium caudatum* ^14, 15^) have elucidated feedback-control mechanisms that tie together three spatiotemporal scales: Fast sub-cellular sensors and actuators operate within tens of milliseconds, leading to cellular reorientation behaviors within about one second, ultimately resulting in directed cell movements and task accomplishment over tens of seconds ^1, 23^. Although many open questions exist, the view has emerged that organismal behaviors are overwhelmingly characterized by the microswimmers rolling around their body axis resulting in helical swimming (hence the term “chiral microswimmer”) ^8,16^, where external stimuli cause either smooth curvature modulation ^3, 16^, or intermittent, randomized body reorientation (“tumble”) ^2, 23^.

In contrast, here we report on a striking microswimmer behavior of *Euglena gracilis* in the form of quantized polygons achieved through highly periodic and regular symmetry breaking of its helical motion (Fig. 1a (i)–(iii), Supplementary Movie S1). Euglena cells achieve these polygonal trajectories by alternating between helical swimming and sharp turning with well-defined lengths and angles, respectively. These polygons typically emerge after a sudden 5-20-fold step-up in light intensity to about ~ 1000 lx. The trajectories are approximately confined to a single plane perpendicular to the light stimulus, and they emerge both near surfaces as well as deep within fluids. Thus, they are not a direct result of interactions with a boundary. We observed polygons with turning angles ranging from 30° to 150°; this angle typically increases over time as the cells adapt to the increased light intensity, yielding polygons of increasing orders, i.e., from order 3 upwards (Fig. 1a (i)–(iii)). These differences from established microswimmer behaviors warrant a deeper investigation of how these polygonal trajectories are generated and what their ethological relevance might be.

**Figure 1:**
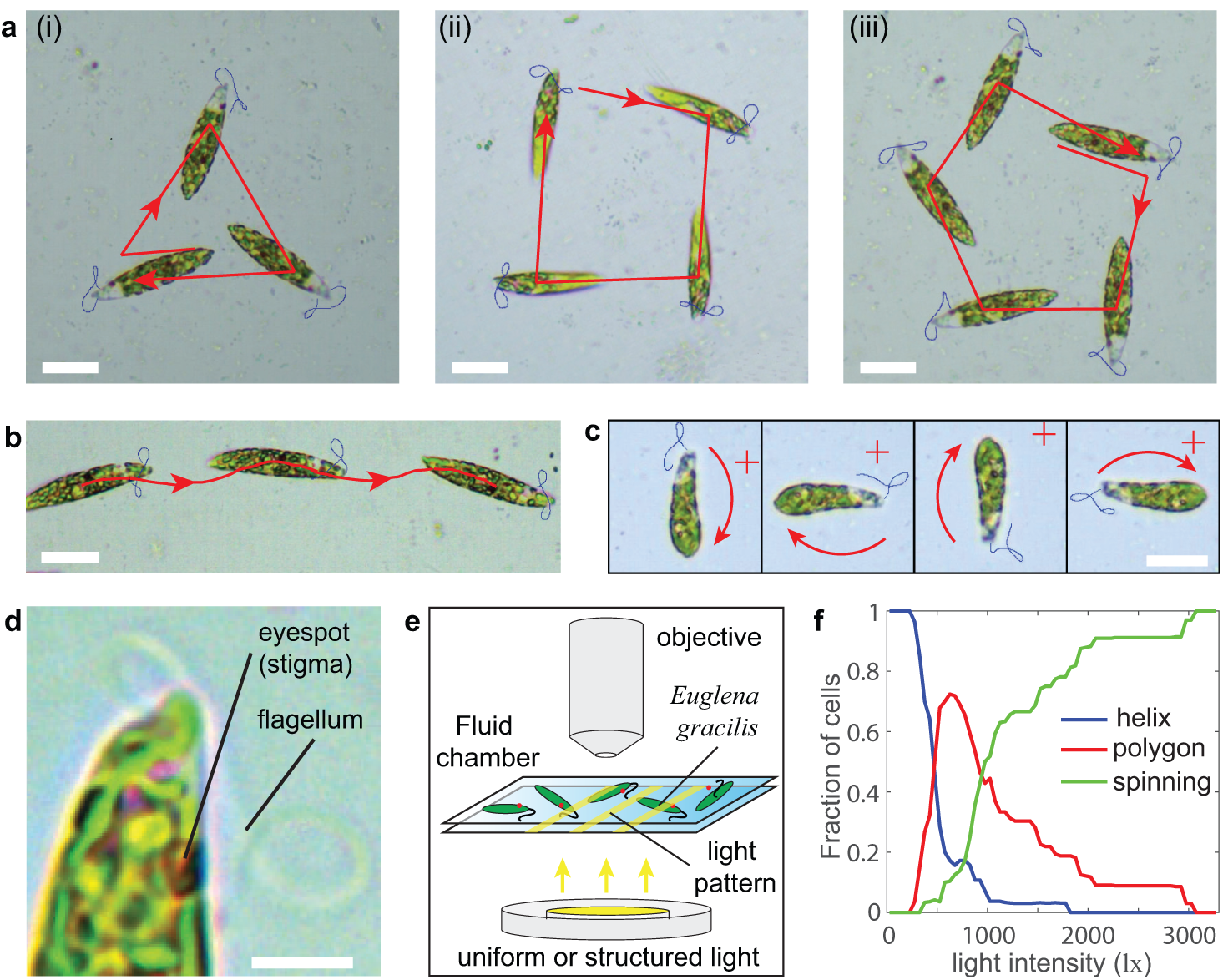
*Euglena gracilis* cells can swim in striking polygonal patterns upon a step-up in light intensity. a, Euglena cells exhibit polygonal swimming trajectories in various orders such as (i) order 3, (ii) order 4, (iii) order 5. These polygonal behaviors are distinct from the previously known behaviors of **b**, helical swimming and **c**, localized spinning. The red “+” symbols mark the same location in space between images. The flagellum outlines are traced and colored in blue. The duration of motion in **a**(i), **a**(ii), **a**(iii), **b** and **c** are 2 s, 2.2 s, 4.1 s, 1.35 s and 0.65 s respectively. **d**, Euglena has an eyespot that shades their photoreceptor. The cell rolls, swims, and turns due to its flagellar beating. This beat is affected by the light intensity sensed by the photoreceptor. **e**, Schematic of experimental setup: the cells are illuminated from below by either uniform or structured light through the microscope lamp or an image projector, respectively. Unless otherwise specified, a uniform light field is applied in the field of view. **f**, We observed different behavioral states of Euglena upon a step-up to various light intensities (always starting from weak light of ~ 50 lx; *n* = 33 cells). The polygonal behavior appears at intermediate light intensities as a transition between helical and spinning behavior. Scale bars: 20 *µ*m (**a**–**c**); 5 *µ*m (**d**).

Euglena is already known to exhibit versatile behaviors, e.g., swimming along helical paths (Fig. 1b, Supplementary Movie S2) at weak light intensities (~ 100 lx) ^6^, or spinning locally (Fig. 1c, Supplementary Movie S3) at much higher light intensities (*>* 3000 lx) ^10^. Euglena has an ellipsoidal cell shape with length of ~ 50 *µ*m and swims at ~ 50–100 *µ*m/s while rolling around its long axis anticlockwise at a frequency of 1–2 Hz ^5, 6^. Its single flagellum beats at ~ 20–40 Hz ^9, 24^, giving 15–20 beats per roll. Euglena senses light signals via a photoreceptor that is partially shaded by the stigma (red ‘eyespot’, Fig. 1d) ^5^. This signal is then converted into different 3D flagellar beating patterns according to the light intensity level ^25^. This response can affect swim speed, rolling frequency, and sideways turning ^5^. These motions then affect the cell’s orientation and position in 3D space, which affects the detected light signal ^6^. This complex feedback enables Euglena to adjust its swimming path in accordance to light conditions to exhibit different phototaxis strategies, e.g., positive and negative phototaxis ^5, 6, 26, 27^ or avoidance turning when encountering a light barrier ^28, 29^.

To better characterize this polygonal swimming phenomenon, we investigated its dependence on light intensity by stepping up the microscope light from ~ 50 lx to a higher intensity (Fig. 1e). We obtained a distribution for the three behavioral states (*n* = 33 cells; Fig. 1f), and we found that the polygonal behavior lies in the transition region between helical swimming and spinning. In the following, we investigate the basis of this polygonal behavior (Fig. 1a), its emergence from the sub-cellular feedback between eyespot and flagellum (Fig. 1d), its relationship to the helical and spinning behaviors (Fig. 1b,c), and the potential biological utility and significance of all three behaviors (Fig. 1a–c), e.g., regarding phototaxis.

First, we investigate the sub-cellular level, to determine what flagellar beat patterns generate polygonal swimming, and how those beat patterns relate to those of helical swimming and spinning (Fig. 2). We manually tracked the flagellum outlines for at least 1.4 s, at a sampling rate of 200 fps (capturing a total of ~ 40 beat cycles, ~ 7 beat patterns per cycle) for 3 cells for each behavior (a total of ~ 2500 frames, Supplementary Fig. S3 and Supplementary Section 3.2). All three behavioral states showed distinct flagellar beat patterns (Fig. 2a–d, Supplementary Movie S4). During helical swimming, the flagellum typically twists into two loops that are distributed on the two sides of the cell (Fig. 2a), whereas for spinning behavior, the flagellum twists into one loop and points to the front of the cell, then subsequently bends to the side opposite to the turning direction (Fig. 2b). Polygonal swimming emerges from periodic switching between two beat patterns that resemble those of helical swimming (Fig. 2c) and spinning (Fig. 2d), which also marks the straight and turning phases of the polygon, respectively. We quantified these different beat patterns with two metrics (Fig. 2e), the maximum vertical distance *f*_1_ from the cell tip and the mean horizontal distance *f*_2_ (defined to be positive on the right of the cell), as well as how the cell orientation *φ* changes over time *t*. For helical beat patterns *f*_1_ remains small (*<* 3 *µ*m), *f*_2_ oscillates around the mean of 0 *µ*m, and *φ* stays nearly constant over time (Fig. 2f). In contrast, spinning beat patterns have large *f*_1_ (~ 6 *µ*m), negative *f*_2_ for all times, and significantly decreasing *φ* over time (Fig. 2g). During polygonal swimming, we find a clear association for *f*_1_, *f*_2_, and the change in *φ* over time (Fig. 2h) with respect to helical and spinning behaviors and the corresponding beat patterns (Fig. 2f,g), respectively. Moreover, we observed that during the turning phase of polygonal swimming, the change in cell orientation |*δϕ*| increases linearly with the number of flagellar beats (a discrete value) (Fig. 2i). Here the slope of |*δϕ*| reveals a turning angle of (18 ± 3)° /beat (always mean ± SEM if not stated otherwise), which varies slightly between different cells, presumably due to the differences in flagellum length and body size. This discreteness is further highlighted by the distinct peaks in the frequency distribution of normalized |*δϕ*| (Supplementary Fig. S4 and Supplementary Section 3.3) for the different beat numbers (Fig. 2j). Thus, we found that the beat patterns for helical swimming and spinning have distinct geometric characteristics, and that polygonal swimming emerges via the switching between these two beat patterns (Fig. 2k,l).

**Figure 2:**
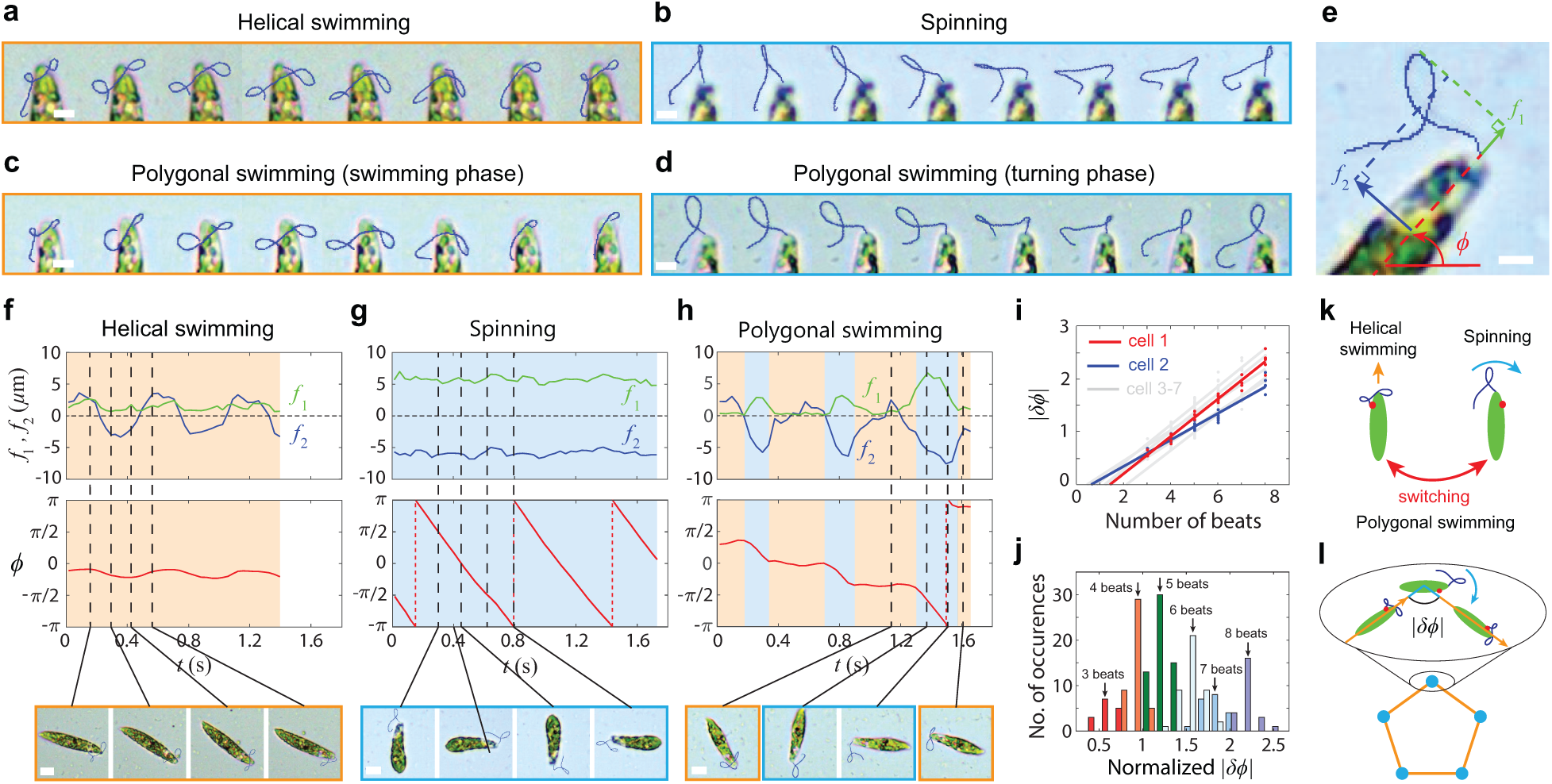
Euglena cells switch between two flagellar beating patterns to achieve the three behavioral states of helical swimming, polygonal swimming, and spinning. **a-d**, Time lapse depicting the representative flagellar beat patterns for each behavior. The flagella are traced in blue. Interval between frames is 200 ms. The helical beat patterns and spinning beat patterns are highlighted by the orange and cyan boxes. **e**, We introduce two metrics, *f*_1_ (green) and *f*_2_ (blue), to characterize the flagellar beat. We also track the cell orientation *φ* as a third variable. **f-h**, Comparing *f*_1_, *f*_2_, and the rate of change of *φ* for all three behaviors reveals that the beat pattern and cell reorientation for polygonal swimming can be characterized as a periodic alternation of helical and spinning behavior. Note that *f*_1_, *f*_2_, and *φ* are obtained from averaging data of flagella outlines in each beat cycle. **i**, The turning angle |*δφ*| during polygonal turning increases in a discrete manner with the number of spinning-like beats. **j**, Histogram for the number of occurrences of different values of normalized |*δφ*| (Supplementary Section 3.3) from 7 different cells, with a total of 202 beat cycles. Each bin has a width of 0.32 and is divided into left and right parts to represent data for a certain number of beats (in different color). For visualization purpose, the plotted width of the bins is reduced and white spaces are added between bins. **k**, Polygonal swimming emerges from the periodically switching between helical and spinning beat patterns. **l**, Periodic turns give rise to a quantizied polygon. Scale bars: 5 *µ*m.

Second, we turned to the cellular level and analyzed the two-way feedback between the 3D cellular reorientation and the light sensed by the eyespot (Fig. 3). The Euglena dynamics can be described with respect to the lab frame and the body frame (Fig. 3a): In the lab frame 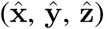, we define the cell orientation *φ* and the phase angle *θ* for the periodic helical trajectory such that there is an apparent angular oscillation of amplitude *ζ* via 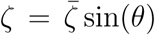 and *θ* lie in the 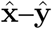 plane, and the light stimulus **I** is parallel to 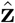. In the body frame 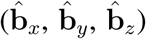, we define 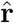 (the direction of maximal light sensitivity, which is assumed to align with the orientation of the eye-spot, i.e., 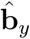; see Supplementary Section 2.1 for a more general model), 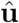 (swimming direction), and 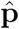 (yitch axis). The cell swims at speed *v* along 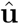, rolls around 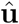 at frequency *ω* (positive for anticlockwise rotation), and turns around 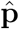. Rather than using the conventional “roll-pitch-yaw” reference system, we substitute the latter two components with a “yitch-paw” vector, 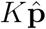. Here the rotations of the body perpendicular to the roll axis are fully described by the “paw-angle” *α* with respect to the eyespot (which defines the unit “paw-vector” 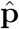 perpendicular to the “roll-axis”) and the ”yitch-rate” *K*, which is the magnitude of rotation around 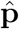.

**Figure 3:**
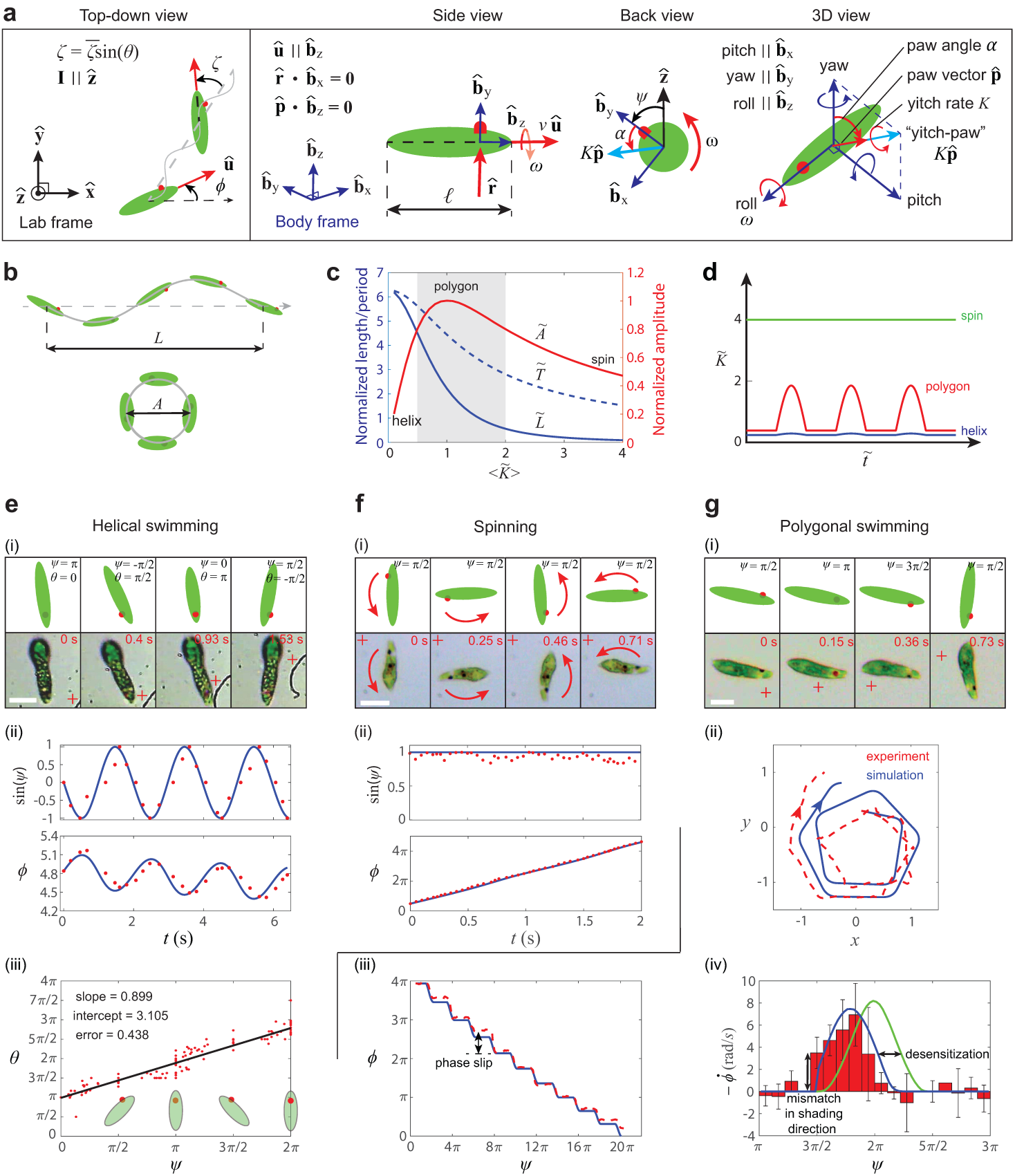
Biophysical model and experiments reveal distinct phase relations between eyespot and cell orientation for the three different behavioral states. **a**, Schematic for definition of model parameters. Light stimulus **I** comes from 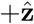 direction. **b**, Side and back view of resulting helical paths. **c**, Relations of *A*, *L* and *T* to 〈*K*〉 (average *K* over the half rolling cycle that **I** is detected), where values of each colored line follow the vertical axis of the same color. The grey region highlights where polygonal swimming occurs. Non-dimensionalized variables are denoted by a “~” on top. **d**, Response functions for different behaviors in response to increase in |**I**| with time. **e**, Helical swimming: (i) Time-lapse depicting eyespot and cell orientation at different phases. (ii) Time evolution of *ψ* and *φ*, experiment (dots) and simulation (lines); (iii) Constant phase relation between *ψ* and *θ*, obtained by linear fit of the experimental data. **f**, Spinning: (i) Eyespot is locked at the same *ψ* during spinning. (ii) Time evolution of *ψ* and *φ*, experiment (dots) and simulation (lines). **g**, Polygonal swimming: (i) Sharp reorientation; (ii) Comparison of path from simulation and experiment. *l* is scaled to 1; (iii) Phase-slip of *φ* with respect to *ψ* due to polygonal turns. (iv) Angular velocity distribution. The error bars denote the standard deviation of experiment data (red bar). The blue and green lines show the mean 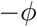 obtained from the simulations. Parameters used in simulations can be found in the Supplementary Section 2.3. For **e**(i), **f**(i), **g**(i), the “+” symbols mark the same point in space between images. Scale bar: 15 *µ*m.

We now consider a generic model where *K* and *α* depend on the light intensity **I**(*t*) detected by the eye spot (while *ω* and *v* are constant). As a first approximation, we assume that the yitch rate *K* depends on the light intensity |**I**| and the orientation of the light sensor relative to the light (i.e., 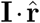, where 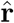 is the sensor’s vector), and is given in terms of *K*_0_, *K*_1_, and *K*_2_ and coupling constants *K_a_* and *K_d_*:

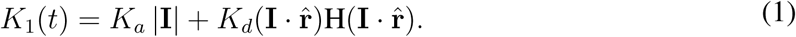

Here *H* is the Heaviside function which accounts for shading by eyespot from one side. The re-ceptor can also exhibit adaptation to light, which is taken into account in *K*_2_(*t*) with the characteristic adaptation rate *γ* (here we assume an adaptation rate much slower than the rolling frequency, i.e., *γ ≪ ω/*(2*π*)):

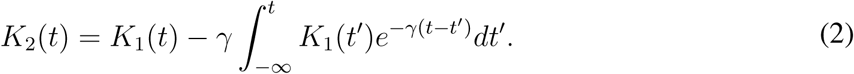

The yitch rate *K* is then given by an intrinsic, light-independent rate *K*_0_ and light-dependent rate *K*_2_:

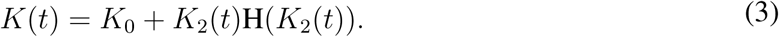

Here the Heaviside function implies that the signal can only be of an activating type. In addition to *K*, the paw angle *α* (Fig. 3a) determines the direction of light-dependent turning, which varies with |**I**| in general, but we assume *α* to be fixed for any given behavior with its value determined by experimental observations.

Simulations of our model successfully capture the transition to polygonal swimming, which occurs between helical swimming and spinning due to the increase in light intensity level (increase in *K*). Here we non-dimensionalize the system by choosing the body length *l* as the length scale, and the inverse of the rolling frequency *ω* as the time scale (Fig. 3a). The helix amplitude *A* first increases with *K*, then decreases at large *K*, while the helix length *L* and the helix period *T* both decrease monotonically with *K* (Fig. 3b,c). Thus, at large *K*, the cell spins at a zero-twist helix with small *A*, *L*, and *T*. Polygonal swimming occurs due to symmetry breaking in light received by the photoreceptor during the rolling cycle: the light is partially blocked by the eyespot, thereby resulting in a “on-and-off” signal in *K* (Fig. 3d). In contrast, in helical swimming and spinning, the cell senses low and high light intensities at all times respectively. These modeling results reveal the importance of the changes in light stimulus sensation due to eyespot rolling in different behavioral states, which we then tested experimentally as described in the following paragraphs.

For weak light intensities 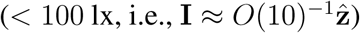, our model predicts that the the frequency of body rolling and helical swimming are coupled, identical, and phase locked (Fig. 3a(i), Supplementary Movie S5). To verify this, we define the roll angle *ψ* using the eyespot as a reference point (Fig. 3a) when observing from the top-down direction 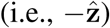. We tracked *ψ*, *φ* and *θ* of the helical swimming cells, and we found a fixed phase relation between *ψ* and *φ* as well as *ψ* and *θ* (Fig. 3e(ii),(iii)), in agreement with our model. By fitting experimental and theoretical results, we found that this phase relation is fixed at *α* = 3.32 ± 0.07 (*n* = 9 cells, each tracked for at least 3 periods), i.e., *α ~ π*. Thus, we conclusively show for the first time that the helical swimming is coupled to the eyespot’s rolling and therefore, the photosensory system.

For strong light intensities 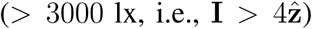, our model predicts spinning behavior. In general, Euglena cells exhibit a variety of complex behaviors at strong light ^10^: They spin locally around their short axis at very high frequency, in either clockwise or anticlockwise direction (we saw no bias in experiments); reverse their rolling and swimming direction; transition to fast helical motion after spinning; or even deform their bodies into rounder conformations. Here we focus only on the spinning behavior, leaving the other behaviors for future study. Our model shows that spinning occurs when the light sensor saturates due to a strong light stimulus (Fig. 3f(i), Supplementary Movie S6). From fitting, we found that *α* = 4.80 ± 0.24 (*n* = 7 cells, over more than 1.5 periods), i.e. *α ≈* 3*π/*2. Thus, spinning results in a yitch rate much larger than the rolling frequency, i.e. *K >> ω*. Furthermore the eyespot stays approximately at the same location with respect to the body, while the orientation varies linearly with time (Fig. 3f(ii)).

For intermediate light intensities 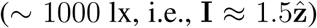, our model predicts polygonal swimming behavior due to periodic shading of the eyespot to the light stimulus (Fig. 3g(i), Supplementary Movie S7), with clockwise (*α ≈ π*) or anticlockwise (*α ≈* 0) trajectories. In our experiments, only clockwise trajectories were observed due to the bias in turning direction resulting from the light coming from +*z* direction. By fitting experimental and theoretical results we found that *α* = 3.18 ± 0.12 (*n* = 9 cells, over more than 3 periods), i.e., *α ≈ π*. Note that we only use a single *α* value which is consistent with helical swimming (see discussion in Supplementary Section 2.1). During polygonal swimming, the cell senses the light stimulus for only half of the rolling cycle due to eyespot shading. The cell then swims in a helix and turns sideways (orthogonal to the light direction) whenever the light is detected. The process then repeats periodically and a polygon emerges. This polygonal path is actually not completely planar but slowly moves along the light direction (approximately 1 body length for 5–10 rolling cycles). The polygonal path tends to increase in size with time, i.e., the order of the polygon increases. This is captured when the simulation accounts for the adaption (Fig. 3g(ii)). For this particular example, we experimentally obtain an adaption time scale of ~ 2 min, beyond which the cell transitions to helical motion. The polygonal turns manifest a “phase-slip” between *ψ* and *φ* (Fig. 3g(iii)). We also observed that Euglena accomplishes the sharp turns much faster than half the roll cycle during which the light sensor is exposed to **I** (red bar from 3*π/*2 to 5*π/*2 in Fig. 3g(iv)), which is not captured by Eqs. (1)–(3) (green line). This difference in time scale can be fixed by a more general model (blue line, Supplementary Section 2.1) that also accounts for the step-up in signal due to a mismatch in sensor’s vector and shading direction as well as the fast signal decay due to short-time desensitization. Thus, the polygonal behavior can indeed be explained by the cell sensing a strong, temporary light stimulus that causes turning, and where the sensor adapts over time.

Third, we investigated the biological utility of these three distinct behaviors on phototactic navigation over longer time and length scales (Fig. 4). We generated different structured light landscapes with an optical projector ^28, 29^ (Fig. 1e, Supplementay Fig. S1 and Supplementary Section 1.3) and observed the cells’ trajectories (Fig. 4a–c, top): (1) When Euglena cells (*n* = 27) coming from low light encounter a step-up to strong light intensity, polygonal swimming or localized spinning occurs, leading to the cells turning around to avoid the light “barrier” (Fig. 4a, Supplementary Movie S8). (2) When suddenly imposing a strong, spatially homogeneous light to cells (*n* = 10) which had previously been at low light, these cells initially spin and then swim in polygons of increasing “search radius” (eventually even exhibiting prolonged phases of helical swimming) before ultimately escaping into a darker region, where they switch back to helical motion (Fig. 4b, Supplementary Movie S9). (3) When suddenly exposing cells (*n* = 15) under low light to a spatial light gradient, they switch between spinning, polygonal swimming, and helical swimming, and essentially execute a biased random walk down the light gradient (reminiscent of bacterial “run-and-tumble” ^2^) (Fig. 4c, Supplementary Movie S10). As the cell travels down the gradient, the variation of 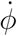 decreases with decreasing light intensity (Supplementary Fig. S5), and over the course of a minute, most cells move away from the lit region (*n* = 15, Supplementary Fig. S6). All these behaviors are also captured in simulations of our model (Fig. 4a–c, bottom, Supplementary Movies S11-13). Thus, coordinated switching between these three cellular behaviors (Fig. 1a–c) enables distinct navigational strategies perpendicular to the direction of light, in addition to the well-described Euglena phototaxis along the light vector ^6^ (which our model also captures but will be detailed in future work).

**Figure 4:**
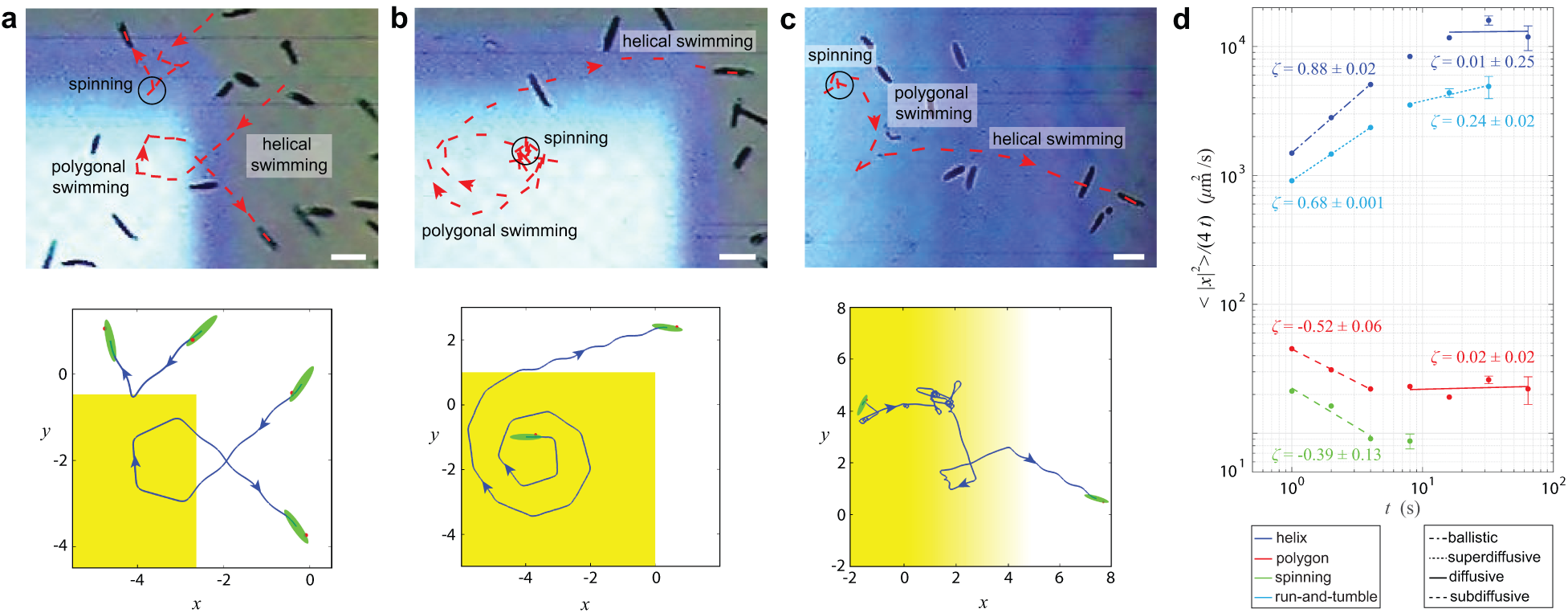
Euglena cells actively tune their anomalous diffusion, exhibiting versatile phototaxis strategies by switching its behavioral state in response to different light structures. **a**, When Euglena cells encounter a sudden step-up in light intensity, they make multiple polygonal turns or directly spin around to avoid this higher intensity light (duration: 5 s). **b**, When Euglena cells are exposed to a sudden increase in light intensity, they initially spin before transitioning to a polygonal path of increasing order, which leads to steadily increasing search radius until the edge of the light field is found and ballistic helical motion sets in (duration: 21 s). **c**, When Euglena cells are exposed to a spatial gradient of light, they switch between spinning, polygonal motion, and helical spinning in a light intensity dependent matter, leading to a biased “run-and-tumble” towards the darker region (duration: 7 s). Bottom rows in **a-c** show the simulations with our model for each of the cases (Supplementary Section 2.3). **d**, Log-log plot of 〈|*x*|^2^〉/(4*t*) of different behavioral states over time *t* (Supplementary Section 3.5). Different forms of anomalous diffusion feature different slopes (i.e, ζ = ~ − 1): subdiffusive (−1 < ζ < 0), diffusive (ζ ~ 0), superdiffusive (ζ > 0), ballistic (ζ ~ 1). The data for helical swimming, polygonal swimming and spinning were collected from traces of 10 cells each. The data for run-and-tumble were collected from 15 traces of the experiment shown in **c**. The error bars denote the SEM and are omitted if smaller than the size of the data point markers. The ± denote the SEM of the piecewise linear fits. Scale bars: 50 *µ*m.

Navigation and search perpendicular to light vectors is relevant for photosynthesis and avoiding UV damage ^5, 30^, e.g., a cell swimming into a bright region from under a leaf (Fig. 4a), or experiencing sudden global changes in sun light intensity due to a cloud (Fig. 4b,c). Although a cell cannot instantaneously discriminate between a spatial or temporal (and global) change in light intensity, it can effectively select the optimal response through adaptation (Fig. 4a-c): It first performs a localized search over the scale of its swim distance in one roll (i.e., 50 − 100 *µ*m) via spinning, followed by a steady increase in search radius through polygonal motion, then intermittently transitions between polygon and helix, ultimately reaching the ballistic motion for the pure helix.

Finally, we quantitatively measured how the switching between these three behavioral states (Fig. 1a– c) (or actually two beat patterns; Fig. 2k) enables the cell to select the magnitude and type of its anomalous diffusion behavior as defined by *〈|x|*^2^*〉* = 4*Dt^∈^*. Here *D* is the generalized diffusion constant and *∈* is the anomalous diffusion exponent ^11, 12^. For convenience, we introduce ζ = ∈ − 1 to represent the slopes of the loglog plot of 〈|*x*|^2^〉/(4*t*) (normal diffusion constant in 2D) of different behavioral states over time *t* (Fig. 4d): (1) Spinning leads to subdiffusive behavior (*ζ ~ -*0.4) as the non-zero forward velocity makes the cell swim in small circles; after *>* 10 s cells often stop spinning. (2) Pure helical swimming leads to ballistic motion with 64 ± 2 *µ*m*/*s (*ζ ~* 0.9, *ζ* is slightly less than 1 due to orientation fluctuation during helical motion), which over longer times transitions to diffusive behavior (*ζ ~* 0) with diffusion constant of *D* = 11000 ± 3000 *µ*m^2^*/*s. (3) Polygonal swimming exhibits subdiffusive behavior at short times (*<* 10 s) due to the looping motion (*ζ ~ −*0.5), while over longer times (*>* 10 s) the cell increases the search radius (Fig. 4b) and transitions to diffusive behavior with (ζ ~ 0) with *D* = 23 ± 2*µ*m^2^/s. (4) For run-and-tumble in the light gradient (Fig. 4c), the cell exhibits superdiffusive behavior due stochastic switching between helical, polygonal, and spinning behaviors, and where scaling decreases over time (*ζ ~* 0.7 for *<* 10 s; and *ζ ~* 0.2 for *>* 10 s); after escape from the light field the motion transitions to scenario (2). Hence the cell can select between ballistic motion and different forms of diffusion (subdiffusive, diffusive and superdiffusive) and tune the corresponding diffusion constant over at least three orders of magnitude (Fig. 4d).

In conclusion, we described a new type of microswimmer behavior which results in polygonal trajectories. This behavior emerges from the light-dependent switching between two beating patterns responsible for helical swimming and localized spinning behaviors, and which are mediated through periodic symmetry breaking as the cell rolls around its axis in the presence of a directional light source. The spinning and polygonal behaviors enable full 2D navigation through translation and turning. Proper timing of behavioral switching with respect to long-axis rolling enables full 3D navigation; however, the fact that the cell trajectory stays approximately within a plane underscores the objective of navigating perpendicular to the light vector. Coordinated, light-intensity dependent switching between both beat patterns allows the cell to actively control and adapt its form of diffusion, enabling tasks like edge detection, local search, and gradient descent in complex light environments. Thus a simple control-feedback loop between cells rolling phase, stimulus detection, and reorientation response binds together multiple system scales (flagellar beats, cellular behaviors, and phototaxis strategies) and automatically tunes itself to the relevant length scale of the light pattern. These results might generalize to behavioral switching in other natural microswimmers and inform the design and control of light-guided synthetic microswimmers ^13, 29^.

## Acknowledgement

We thank members of the Riedel-Kruse lab, N. Ouellette, and A. Macdonald. This work was supported by NSF #1324753 and the Croucher Foundation (through a postdoctoral fellowship to A.C.H.T.) Author Contributions: Project Idea: ACHT and IHRK; Theory: ACHT and IHRK; Modeling: ACHT; Experiments and Data Analysis: ACHT (except Fig.4: ACHT and ATL); Manuscript Preparation: ACHT and IHRK.

